# Guppies prefer to follow large (robot) leaders irrespective of own size

**DOI:** 10.1101/320911

**Authors:** David Bierbach, Hauke J. Mönck, Juliane Lukas, Marie Habedank, Pawel Romanczuk, Tim Landgraf, Jens Krause

## Abstract

Body size is often assumed to determine how successful an individual can lead others with larger individuals being more likely to lead than smaller ones. However, direct evidence for such a relation is scarce. Furthermore, even if larger individuals are more likely to lead, body size correlates often with specific behavioral patterns (e.g., swimming capacity) and it is thus unclear whether larger individuals are more often followed than smaller ones because they are larger or because they behave in a certain way. To control for behavioral differences among differentially-sized leaders, we used biomimetic robotic fish – Robofish – of different sizes. Robofish is accepted as a conspecific by live guppies (*Poecilia reticulata*) and provides standardized behaviors irrespective of its size. We specifically asked whether larger leaders are preferentially followed when behavior is controlled for and whether the preferences of followers depend on their own body size or their risk taking behavior (‘boldness’). We found that live guppies followed larger Robofish leaders closer than smaller ones and this pattern was independent of the followers’ own body size as well as risk-taking behavior. This is the first study that shows a ‘bigger is better’ pattern in leadership in shoaling fish that is fully independent of behavioral differences between differentially-sized leaders and followers’ own size and personality.

## Introduction

What defines how successfully an individual can lead others? In shoaling fish, those individuals that occupy front or periphery positions within a shoal are assumed to have the greatest influence on the group’s movement direction, hence are capable of leading the other shoal members [1–4]. Often, occupation of front or peripheral positions is related to motivational or phenotypical differences among individuals [2, 5]. For example, individuals that go front are hungrier [6, 7], more risk-taking (‘bolder’) [4, 8–11] or simply larger [5, 12] than the rest of the shoal. Mechanistically, those individuals at front may swim faster [4, 12] or have larger repulsion areas [5, 13], both leading to an assortment within the shoal. However, being at the front (i.e., taking the lead) is often not the only factor determining leadership success. Using the golden shiner (*Notemigonus crysoleucas*), Reebs ([14]) showed that a minority of informed large fish was capable of leading a shoal of small fish to a food location but informed small fish had much lower success in leading a shoal of large fish even when occupying the front positions in the shoal. Furthermore, when sticklebacks (*Gasterosteus aculeatus*) were grouped with two partners of different personalities, they were more likely to follow the partner of similar personality out of cover [8]. Thus, both body size as well as behavior may determine leadership success in fishes. In addition, both body size and behavior often covary with each other, for example larger fish can swim faster than small ones [1] or exhibit a certain personality [15] and only recently Romenskyy et al. [13] concluded that “fish of different sizes cannot be considered simply as particles of different physical size, since their behavior changes with their size”. It is thus unclear whether larger individuals are more often followed than smaller ones because they are larger or because they behave in a certain way. Furthermore, we do not know whether following depends on the followers’ own size or behavior and how follower size and behavior may interact with leader size. To answer these questions it is necessary to experimentally control for the leader’s behavior while simultaneously varying its body size (or vice versa).

In the current study, we used a biomimetic robotic fish – Robofish – that is accepted as a conspecific by live fish (guppy, *Poecilia reticulata*, [16, 17]) to gain control over the behavior of the leader. We asked (a) whether larger leaders are preferentially followed (as predicted by a “bigger is better” hypothesis) when behavior is controlled for and (b) whether the preferences of followers depend on their own body size or their risk taking behavior (‘boldness’).

## Methods

### Study organism and maintenance

We used Trinidadian guppies (*Poecilia reticulata*) that were descendants of wild-caught fish from the Arima River in North Trinidad. Test fish came from large, randomly outbred single-species stocks maintained at the animal care facilities at the Faculty of Life Sciences, Humboldt University of Berlin. We provided a natural 12:12h light:dark regime and maintained water temperature at 26°C. Fish were fed twice daily *ad libitum* with commercially available flake food (TetraMin™) and once a week with frozen *Artemia* shrimps.

### The Robofish system

The Robofish is a three-dimensional (3D)-printed guppy-like replica that is attached to a magnetic base. The magnetic base aligns with a wheeled robot that is driving below the actual test tank (88 × 88 cm) on a transparent second level. Hence the replica can be moved directly by the robot (Figure 1). The entire system is enclosed in a black, opaque canvas to minimize exposure to external disturbances. The tank is illuminated from above with artificial light reproducing the daylight spectrum. On the floor, a camera is facing upwards to track the robot’s movements through the transparent second level. A second camera is fixed above the tank to track both live fish and replicas. Two computers are used for system operation: one PC tracks the robot, computes and sends motion commands to the robot over a wireless channel. The second PC records the video feed of the second camera which is afterwards tracked by custom-made software [18]

**Figure 1:**
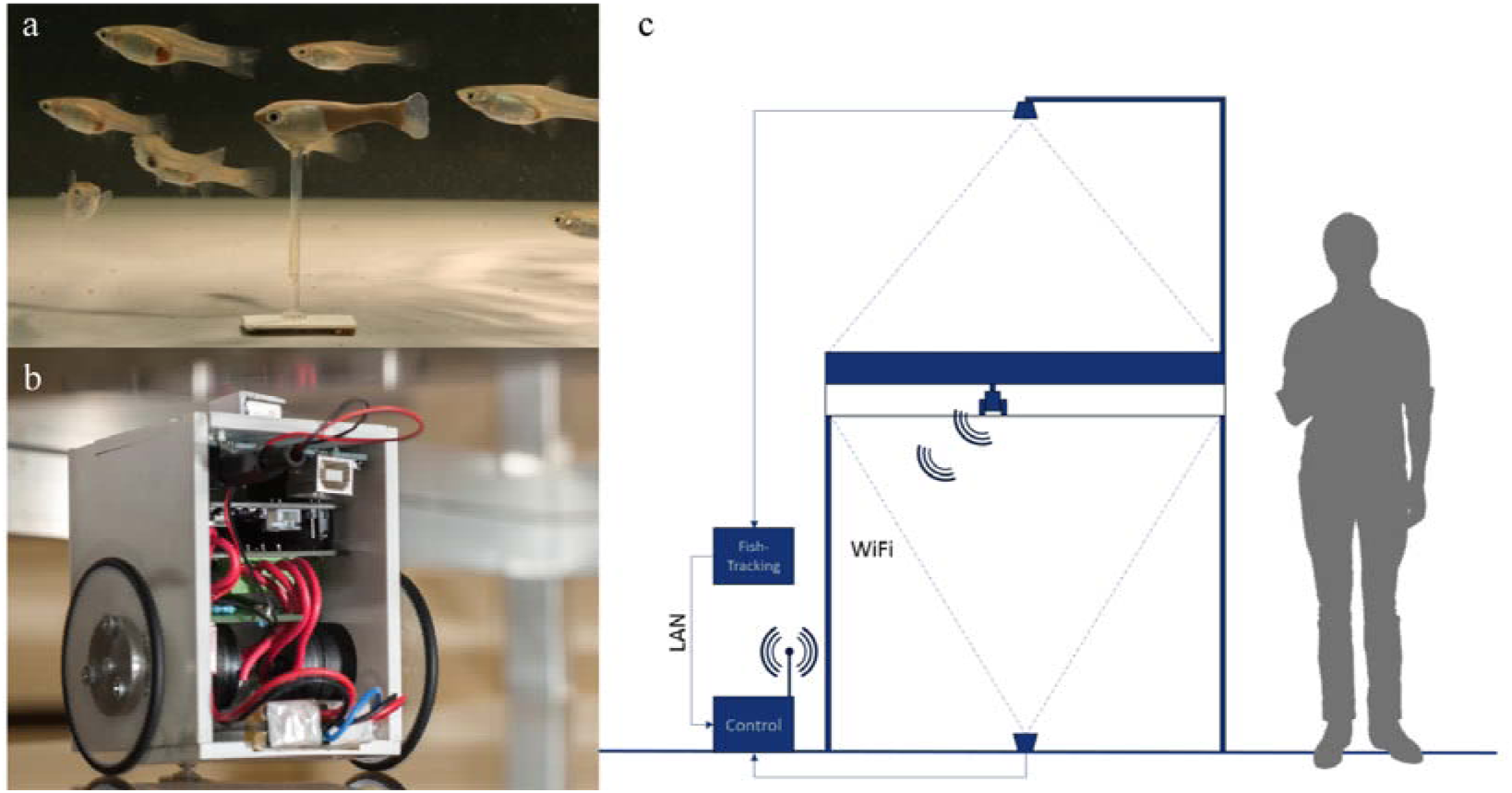
The Robofish system. (a) Guppy-like replica (3D printed and colored) with a group of female guppies in the test arena. (b) Close-up of the robot unit. The robot unit is driving on a second level below the test arena (c).

### Experimental setup

To provide live guppies with differently-sized Robofish leaders, we used three replicas that differed only in body size (r1=20 mm standard length (SL); r2=25 mm SL, r3=30 mm SL, see Figure S1). As we used transparent screws to attach the replica to its magnet foot, all replicas regardless of size kept the same distance to the water surface (1 cm, at 10 cm water level). Similarly, we used live test fish (only females to avoid sex effects) that were preselected into three body size classes: 20 mm (ranging from 18.0 to 21.9, mean=20.1, *n*= 24), 25 mm [23.7 to 27.2, mean= 25.2 *n*= 33], 30 mm [28.0 to 32.0, mean= 30.0 *n*=33]. Thus, we had a balanced two-way design with the factors leader size (three levels) and live fish size (three levels).

To initiate a trial, we transferred our test fish into an opaque PVC cylinder located at the lower left corner of the test tank. The PVC cylinder had an opening (diameter 3 cm) which was closed with a sponge. We removed the sponge after 1 minute of acclimation and we noted the time each fish took to leave the cylinder as a proxy for its risk taking tendency (‘boldness’) which might correlate with following tendencies [10, 19, 20]. We started the Robofish’s movement when the live fish left the cylinder (= one body length away from the cylinder’s border). The Robofish moved along a zigzag path to the opposite corner and then counter-clockwise to its start position (see Figure 2 for an example track as well as video in the supplement SI_video_1). This round was repeated for a second time and a trial took about 60 s in total. Each trial was videotaped for subsequent tracking and the test fish was transferred back to its holding tank. Based on the tracked positions, we calculated the inter-individual distance (IID) between focal fish and Robofish, which has been shown to reflect a live fish’s tendency to follow the moving Robofish [20].

**Figure 2:**
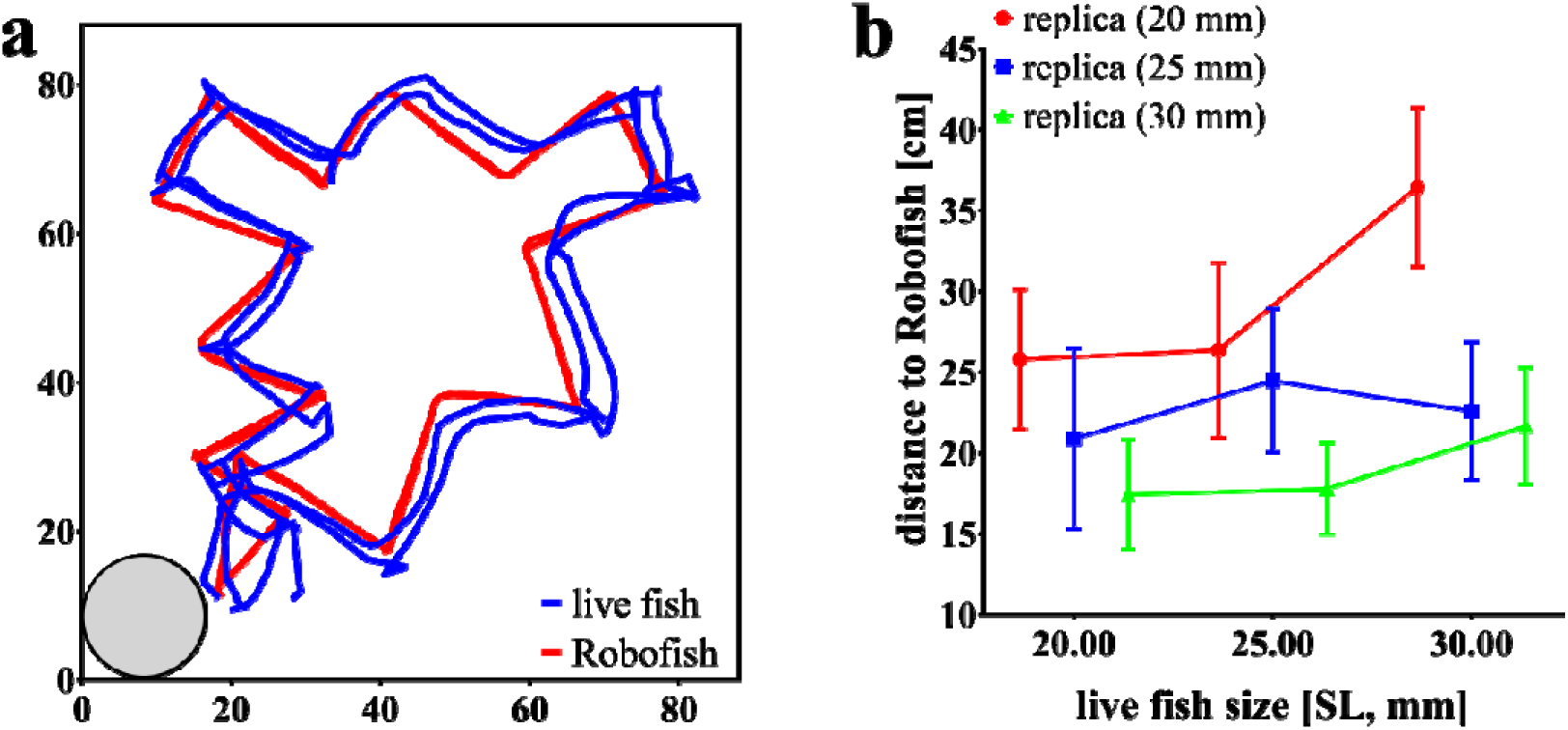
Following behavior of live guppies towards differently sized Robofish replicas. (a) Example track of a trial with Robofish. Fish were introduced into the start box (grey circle, lower left corner) and released into the tank after 1 minute. Robofish then moved on a predefined zig-zag trajectory through the tank until it reached its start position. This movement was repeated a second time and a trial lasted about 60s in total. (b) Inter-individual distance (IID) between live fish and Robofish replicas. Shown are means (± S.E.M) for each size combination.

### Statistical analysis

We initially log10 transformed both recorded continuous variables (IID, time to leave start box) to match a Gaussian distribution. We then used the IID as dependent variable in a ANOVA (unianova package in SPSS 25) with ‘leader size’, ‘live fish size’ and their interaction term as fixed factors and ‘time to leave start box (log10)’ as covariate. In order to test whether differently sized live fish differ in their risk aversion tendency, we further correlated live fish body size with time to leave shelter using Spearman’s rank order correlation.

## Results

Regardless of own size (non-significant factor ‘live fish size’ *F*_2,78_= 1.52; *p*=0.23), live guppies followed larger Robofish replicas significantly closer than smaller ones (significant effect of factor ‘leader size’ *F*_2,78_= 4.49; *p*=0.014, figure 2). There was no size assortative pattern detectable (i.e., smaller live fish did not follow smaller replicas closer than larger ones and *vice versa*) as suggested by a non-significant interaction term ‘leader size × live fish size’ (*F*_4,78_= 0.49; *p*=0.74). Also, the time each fish took to leave the start box had no significant influence on its following behavior (*F*_1,78_=0.90; *p*=0.35).

There was no significant correlation between live fish’s body size and their tendency to leave the start box (Spearman’s *r*=0.186, *p*=0.08).

## Discussion

Live guppies followed larger Robofish leaders closer than smaller ones and this pattern was independent of the followers’ own body size as well as risk-taking behavior. While keeping the leaders’ behavior constant through the use of a biomimetic robot, this is the first study that showed a ‘bigger is better’ pattern in leadership in shoaling fish that is fully independent of behavioral differences between differentially-sized leaders.

Body size is often inevitably linked to specific behavioral patterns [15] and it is thus experimentally difficult to disentangle what cue (body size or linked behavior) is used by individuals that have to choose among differently-sized conspecifics. While researchers from the field of sexual selection make use of video animations in binary choice tests to decouple behavior from body size and keep either one constant while varying the other (see [21–24]), the study of collective movement has largely relied on the use of live stimuli (but see [25] for a working Virtual Reality set-up). We addressed this issue by using a biomimetic robot that is accepted by live fish as a conspecific and provided the same behavioral cues while varying only body size. With this technique, we show that the leading success of larger individuals is determined by their large body size and does not require differences in swimming speed, boldness or occupied spatial position.

It was previously found that large guppy females prefer to shoal with similar-sized (large) but not smaller conspecifics while small guppy females did not shoal assortatively but also preferred to shoal with larger females [26]. Small individuals can benefit from associating with larger conspecifics if the larger ones take away the attention of harassing males [27, 28] or predators [29]. Large individuals can benefit from associating with other large individuals to minimize the oddity effect during predation [30]. However, also indirect benefits might play a role as larger body size might often be an indicator of fitness (longer survival, better foraging abilities, higher dominance rank) and it thus might be beneficial for followers (regardless of own size) to associate with those successful phenotypes.

We do not know whether live fish may have an intrinsic preference for large body size (as assumed in mate choice contexts, [31]) or larger leaders are simply better visible than smaller ones and thus elicit stronger (retinal) stimulation which is translated into a stronger following response (see [32] for an example of a visual field reconstruction in shoaling fish).

In contrast to the study by Reebs [14] we found no evidence that larger individuals follow less than smaller ones. This can be due to species-specific difference as Reebs [14] used the obligate shoaling golden shiner while we used the facultative shoaling guppy. As shoal membership in fishes is highly dynamic and individuals may maximize their fitness by switching frequently between groups of varying size and composition in response to changes in their physiological stage and the external environment [33], we argue that following behavior can be indeed independent of own size [2]. Also, we found no evidence that follower’s risk-taking behavior affected their tendencies to follow differentially-sized leaders. This result is in contrast to studies on sticklebacks where shyer individuals are better followers and are less likely to initiate leading themselves [8]. Besides possible species-specific differences, we argue that reinforcing feedbacks due to mutual influences among live animals may have led to the observed personality-dependent following behavior in sticklebacks [8, 34].

Our study shows a preference of shoaling fish to follow larger over smaller leaders. We argue that fish, irrespective of their own size have an inherent preference to follow larger leaders, as doing so provides either benefit for the follower or larger leaders are more visible and thus easier to follow.

## Acknowledgments

Experiments reported in this study were approved by the LaGeSo Berlin under the registration number 0117/16. We would like to thank David Lewis for his valuable help in raising our test fish. Furthermore, we like to thank Angelika Szengel and Hai Nguyen for their help in developing the Robofish. We received financial support by the DFG (BI 1828/2-1, RO 4766/2-1, LA 3534/1-1). The authors declare no competing interests.

## Supplemental Information

**Figure SI:**
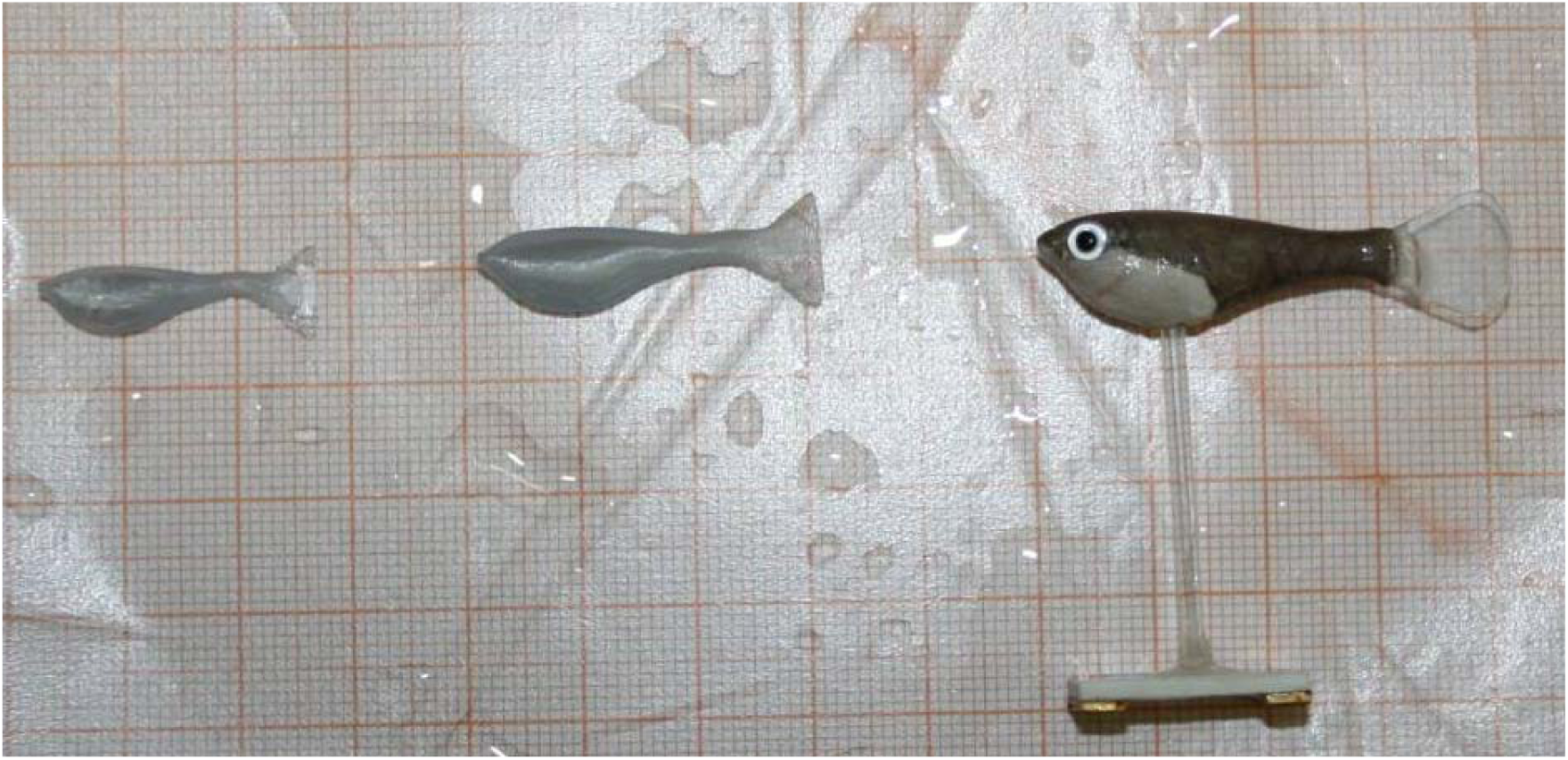
Photograph of differently-sized replicas. Left (20 mm SL) and middle (25 mm SL) replicas are unprocessed 3D printed blanks that were latter on equipped with glass eyes and color-painted as shown for the 30 mm replica on the right.

**Video SI:** Example of a 30 mm live guppy that follows a 25 mm Robofish replica. The live fish leaves the cylinder at 01:04 min.

